# SparK: A Publication-quality NGS Visualization Tool

**DOI:** 10.1101/845529

**Authors:** Stefan Kurtenbach, J. William Harbour

## Abstract

While there are sophisticated resources available for displaying NGS data, including the Integrative Genomics Viewer (IGV) and the UCSC genome browser, exporting regions and assembling figures for publication remains challenging. In particular, customizing track appearance and overlaying track replicates is a manual and time-consuming process. Here, we present SparK, a tool which auto-generates publication-ready, high-resolution, true vector graphic figures from any NGS-based tracks, including RNA-seq, ChIP-seq, and ATAC-seq. Novel functions of SparK include averaging of replicates, plotting standard deviation tracks, and highlighting significantly changed areas. SparK is written in Python 3, making it executable on any major OS platform. Using command line prompts to generate figures allows later changes to be made very easy. For instance, if the genomic region of the plot needs to be changed, or tracks need to be added or removed, the figure can easily be re-generated within seconds without the manual process of re-exporting and re-assembling everything. After plotting with SparK, changes to the output SVG vector graphic files are simple to make, including text, lines, and colors. SparK is publicly available on GitHub: https://github.com/harbourlab/SparK.

## Introduction

Next generation sequencing (NGS) data is usually visualized using tools like IGV or the UCSC genome browser, allowing quick visualization of genomic regions of interest (1,2). Both tools are standard bioinformatic resources that offer considerable functionality and exporting functions. However, manually editing output files into publication-quality figures, exchanging tracks, changing the region of plotted genomic region, and plotting replicate data is difficult and time consuming. As a result, replicate data are usually plotted separately to show all tracks, resulting in visually complex figures that can be difficult to interpret. Consequently, many publications display only one replicate, with the second being supplied as a supplemental file, often in formats that are not easily accessible. Here, we present SparK, a Python 3 program that generates publication quality figures for defined genomic regions, allows tracks to be overlaid, standard deviations to be plotted, and replicate NGS data to be plotted in a visually intuitive manner. Using command line prompts, figures can be re-generated within seconds.

## Results

The following figures are examples of plots generated with SparK. These plots are auto-generated, without any further manipulation. Figure 1 shows the most basic way of plotting NGS data with SparK. In this example, four ChIP-seq tracks are depicted. Standard plotting includes autoscaling of tracks by default, displaying genes, and a scale bar, which is automatically adjusted to be of meaningful size and value, regardless of the size of the plotted region (See Figures 1-4). Transcription start sites are marked with arrows, indicating the direction of transcription. By default, SparK will merge all known transcripts of individual genes in the region, and indicate the first TSS site on the merged gene. Individual transcripts may be plotted instead, as described on the GitHub page. SparK offers different options to visualize replicate data. Figure 2A showcases two features of SparK. First, SparK can overlay tracks by making them semi-transparent.

**Figure 1:**
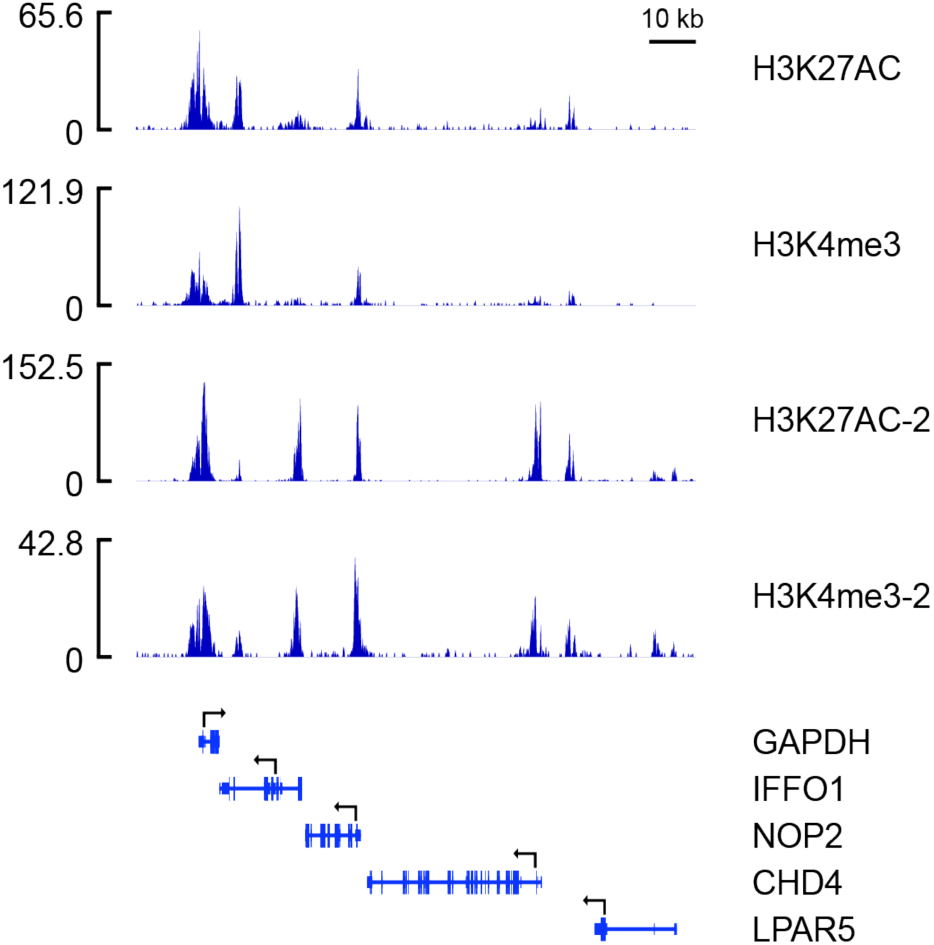
Basic SparK figure plotting four ChIP-seq tracks, gene annotations, and a scale bar.

**Figure 2.**
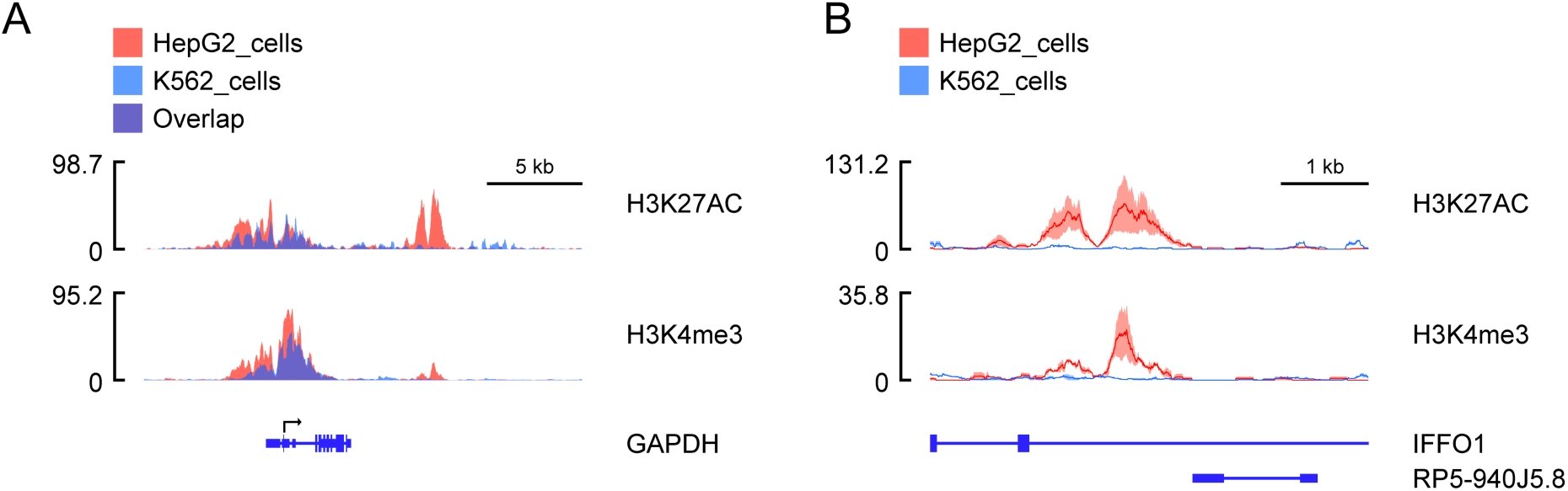
Options of plotting replicates with SparK. (A) Averaged and overlaid replicates of ChIP-seq data derived from two cell lines. (B) Standard deviation plot of the same data, zoomed in on the significantly changed peak downstream of GAPDH. The average of all replicates is drawn as a line, and the standard deviation indicated in the respective color.

**Figure 3:**
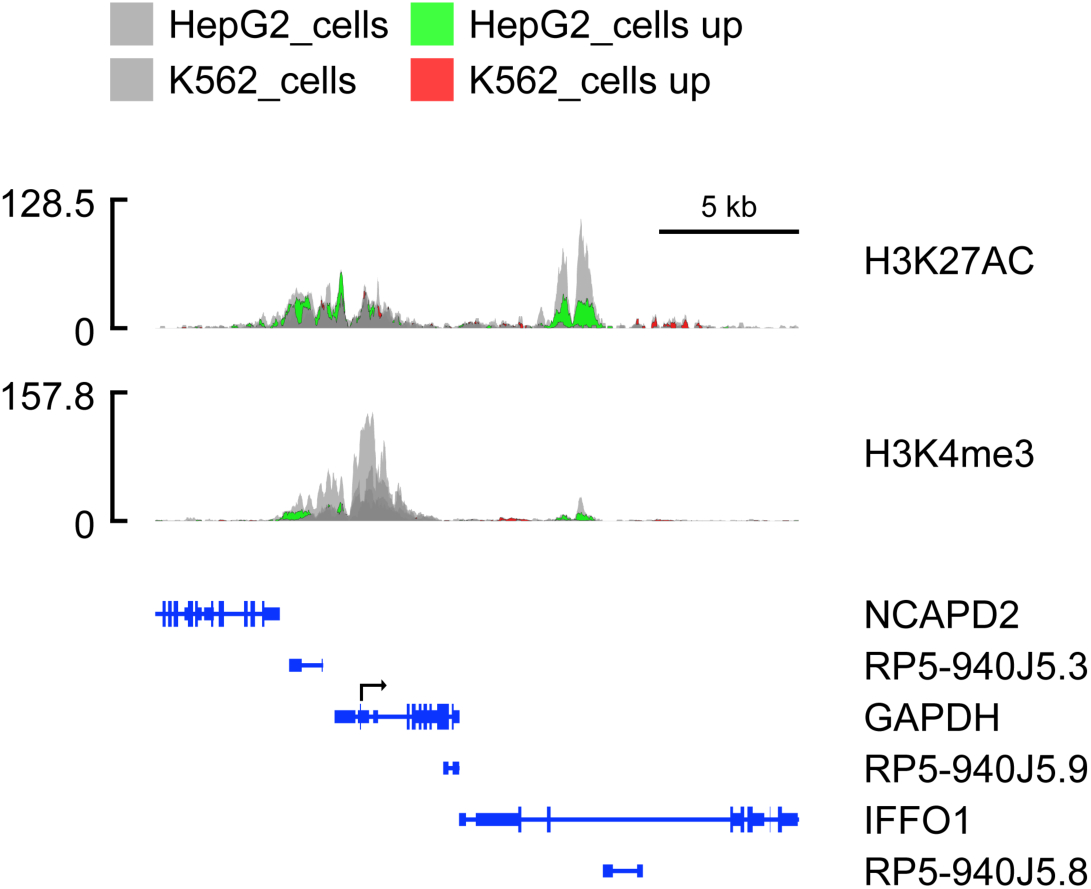
Plots indicating areas with significant changes between two cell lines. Areas (or sparks), are highlighted with green and red.

**Figure 4:**
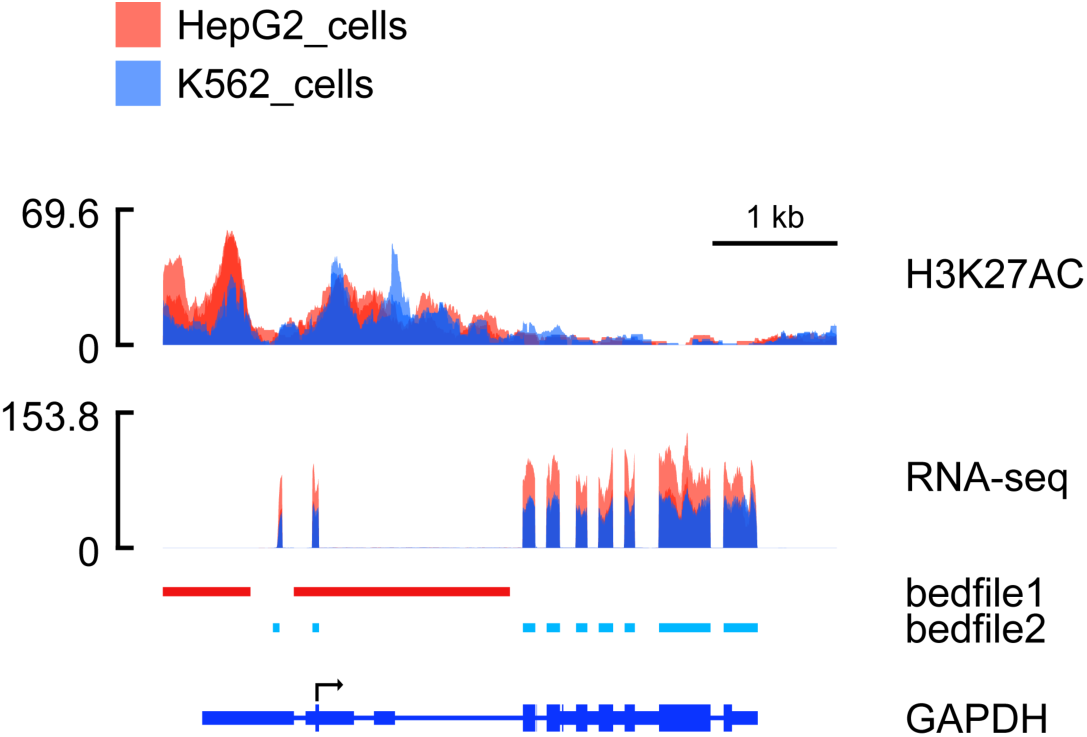
Example plot of mixed datatypes. Two replicates are plotted for each track.

Second, SparK can plot average tracks, rather than displaying all replicates, which can make the plot difficult to interpret. Figure 2B shows the ability of SparK to plot standard deviation tracks, thereby allowing this statistical information to be visualized simultaneously from all replicates.

SparK further offers the option to visualize differences between two groups using “sparks,” which are defined as areas where the difference between the means of the two groups is greater than the sum of their standard deviations. In Figure 3, all replicates were plotted in grey, with sparks indicating differences between the two cell lines. Figure 4 shows the ability of SparK to visualize mixed datatypes. In this example, ChIP-seq and RNA-seq data are displayed along with two bed files with custom coloring. Two replicates for each NGS are plotted for both cell lines. As this case shows, plotting eight NGS tracks in two rows can yield a very useful representation of the data. For simpler visualization, replicate data can be averaged, as was shown in Figure 2A.

## Discussion

Here we present SparK, an easy-to-use Python program to plot NGS data. SparK is a specialized plotting program to generate highly editable and publication quality figures, downstream of existing and powerful analysis tools like IGV and the UCSC genome browser. Besides the functionality described here, SparK offers several customization options that are outlined on the GitHub page.

